# Epigenetic clocks for mice based on age-associated regions that are conserved between mouse strains and human

**DOI:** 10.1101/2022.03.23.485470

**Authors:** Juan Perez-Correa, Vithurithra Tharmapalan, Hartmut Geiger, Wolfgang Wagner

**Affiliations:** Institute for Stem Cell Biology and Cellular Engineering, RWTH Aachen University Medical School, Aachen, Germany; Helmholtz Institute for Biomedical Engineering, RWTH Aachen University Medical School, Aachen, Germany; Institute of Molecular Medicine, Ulm University, Ulm, Germany

**Keywords:** aging, methylation, epigenetics, homology, conserved, predictor, mice, human

## Abstract

Aging of mice can be tracked by DNA methylation changes at specific sites in the genome. In this study, we used the recently released Infinium Mouse Methylation BeadChip to compare such epigenetic modifications in C57BL/6 (B6) and DBA/2J (DBA) mice. We observed marked differences in age-associated DNA methylation in these commonly used inbred mouse strains, indicating that epigenetic clocks for one strain cannot be simply applied to other strains without further verification. In B6 mice age-associated hypomethylation prevailed with focused hypermethylation at CpG islands, whereas in DBA mice CpG islands revealed rather hypomethylation upon aging. Interestingly, the CpGs with highest age-correlation were still overlapping in B6 and DBA mice and included the genes *Hsf4*, *Prima1*, *Aspa*, and *Wnt3a*. Notably, *Hsf4* and *Prima1* were also top candidates in previous studies based on whole genome deep sequencing approaches. Furthermore, *Hsf4*, *Aspa*, and *Wnt3a* revealed highly significant age-associated DNA methylation in the homologous regions in human. Subsequently, we used pyrosequencing of the four relevant regions to establish a targeted epigenetic clock that provided very high correlation with chronological age in independent cohorts of B6 (R^2^ = 0.98) and DBA (R^2^ = 0.91). Taken together, the methylome differs extensively between B6 and DBA mice, while prominent age-associated changes are conserved among these strains and even in humans. Our new targeted epigenetic clock with 4 CpGs provides a versatile tool for other researchers analyzing aging in mice.

## Introduction

Precise measurement of aging is a prerequisite to identify parameters that may attenuate the aging process. It is fascinating that the DNA methylation (DNAm) patterns change in a highly reproducible and seemingly organized manner during aging of the organism (Fraga et al., 2005; Orozco et al., 2014). This epigenetic modification at the cytosine residues of CG dinucleotides (CpGs) impacts on chromatin organization, transcription factor binding, and gene expression. It is therefore anticipated that age-associated DNAm might be of immediate functional relevance for the aging process, albeit this remains to be proven. Today, epigenetic clocks are considered to be the most accurate biomarker for age predictions and there is sound evidence that they also capture aspects of biological aging that are independent from chronological age (Bell et al., 2019; Marioni et al., 2015).

More than ten years ago, the first epigenetic aging signatures have been identified for humans using Infinium BeadChip methylation datasets (Bocklandt et al., 2011; Koch and Wagner, 2011). This microarray platform provides DNAm levels at single CpG resolution (Bibikova et al., 2011) - first on the 27k BeadChip for about 27,000 CpGs, then with the 450k BeadChips for 450,000 CpGs, and currently with the EPIC BeadChips for 850,000 CpGs. With the advent of a rapidly growing number of such datasets, and with improved bioinformatics approaches, epigenetic age-predictors that correlated very well with chronological age were developed for humans (Hannum et al., 2013; Horvath, 2013; Weidner et al., 2014). The development of the bead-array based aging clocks for mice was somewhat trailing the development of human clocks, because BeadChip platforms were initially not available for non-human species (Gujar et al., 2018; Wagner, 2017). Only recently, new tools became available that made the BeadChip technology also applicable to other mammals, such as the Mammalian Methylation array that can measure about 36,000 highly conserved CpGs across mammalian species (Arneson et al., 2022). Furthermore, Illumina has recently released a Mouse Methylation BeadChip for measurement of more than 285,000 CpGs across the mouse genome. So far, research with this new microarray platform with respect to aging clocks has not yet been published.

Several other methods are available to analyze whole genome DNAm patterns (Blueprint-consortium, 2016). Whole genome bisulfite sequencing (WGBS) might give the most comprehensive insight, since it addresses theoretically all CpGs in the genome. However, due to the limited sequencing coverage for specific CpG sites, analysis of age associated CpG sites may be difficult (Zhou et al., 2019). In addition, the method is costly and integration of many datasets remains a challenge. Reduced representation bisulfite sequencing (RRBS) enriches for areas of the genome with a high CpG content to reduce the costs but not all CpGs are analyzed across the samples. Since the Infinium BeadChip technology was not available for non-human species the first epigenetic clocks for mice and other mammals were initially derived from WGBS and RRBS data (Petkovich et al., 2017; Stubbs et al., 2017; Wang et al., 2017). Based on these sequencing profiles, we have previously selected three age-associated CpGs in the genes *Prima1*, *Hsf4*, and *Kcns1* for targeted DNAm analysis with pyrosequencing to facilitate precise estimation of chronological age in murine blood samples (Han et al., 2018; Han et al., 2020). This targeted signature indicated that epigenetic aging is accelerated in the shorter-lived DBA/2J (DBA) as compared to the commonly used C57BL/6 (B6) mice. A systematic comparison how age-associated changes in DNAm patterns vary between these inbred mouse strains is not yet available.

In this study, we utilized the Mouse Methylation BeadChip to compare age-associated DNAm profiles in blood cells from B6 or DBA animals. The age-related changes varied significantly between these strains, which might partly be attributed to general differences of their methylome. Despite these differences, there was also a large overlap of age-associated DNAm that was then used to train an epigenetic clock for the Mouse Methylation BeadChip with 105 CpGs. Notably, several of the top candidate CpGs revealed also age-associated DNAm in corresponding human genomic regions. Based on these results, we introduce a further optimized and thus more reliable 4 CpG pyrosequencing-based epigenetic clock for mice.

## Materials and methods

### Murine blood samples

Blood samples from DBA/2J and C57BL/6 mice were collected at the University of Ulm by submandibular bleeding (100–200 μl) of living mice or postmortem from the vena cava or from the heart, for both training and validation sets. All mice were accommodated under pathogen-free conditions. The experiments were approved by the Regierungspräsidium Tübingen.

### Mouse Methylation BeadChip and data processing

Genomic DNA was isolated using a Qiagen QIAamp DNA Mini and Blood Mini Kit (Qiagen, Hilden, Germany) and measured with a NanoDrop2000 spectrophotometer (Thermo Scientific, Wilmington, USA). 500 ng per sample of isolated DNA was bisulfite converted and hybridized with the Infinium Mouse Methylation BeadChip (Illumina Inc., San Diego, CA, USA), from 24 C57BL/6 mice (all female) and 12 DBA/2J mice (6 male, 6 female). The platform interrogates more than 285,000 methylation sites per sample at single-nucleotide resolution. The association to CpG islands, shelve, and shore regions was taken from the Illumina BeadChip annotation. Raw data from IDAT files was read and processed in R with the ENmix package, and the beta values were normalized using the ENmixD method (ENmix background correction and RELIC dye-bias normalization)(Xu et al., 2016). Probes with detection p-values > 0.01 or more than 10% of NA values were filtered out. One sample with more than 10% missing values was not considered for further analysis. Furthermore, CpGs on X and Y chromosomes were not considered for epigenetic clocks. Probe-type bias adjustment was performed with the Regression on Correlated Probes method (Niu et al., 2016). Initially, quantile normalization for ENmixD was included, which provided a similar selection of age-associated CpGs. However, since this normalization regimen masked the general shifts in age-associated DNAm, we pursued without quantile normalization. To identify significantly differentially methylated CpGs between B6 and DBA mice, the Wilcoxon rank-sum test with Bonferroni-correction (p < 0.01) for the total amount of CpGs in the Mouse Methylation BeadChip was used, after removing the probes in X and Y chromosomes, cross-reactive probes, and probes with single nucleotide polymorphism (SNP).

### Selection of age-associated CpGs and derivation of the 105 CpG predictor

In this study, the initial selection of relevant CpGs was based on the slope of linear regressions of DNA methylation and chronological age to filter for CpGs that have large changes in absolute DNAm levels. Since the two mouse strains have different life expectancy, we normalized the data according to the sex and strain of the mice (Yuan et al., 2009). The CpGs were then filtered independently for B6 and DBA mice by the slope (greater than 30%). The slope of the correlation for each CpG was calculated:

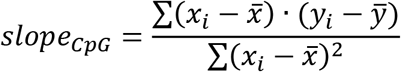

Where *x*_*i*_ is the normalized age of the sample *i*, 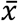 is the mean normalized age of the samples, *y*_*i*_ is the normalized beta value of the sample *i* for the CpG, and 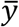 is the mean of the normalized beta values of the CpG. The overlap of the CpGs with slopes >30% in both mouse strains was 105 CpGs. Since the sample number was not sufficient to generate a multivariable model for this relatively large signature we used age-estimations for the individual CpGs with linear models using the normalized beta values (slope and intercept with age). The average of all these single-CpG estimations was considered as the final age prediction.

### Identification of human homolog regions

Homology alignments of human and murine genomes for the 105 CpGs were performed with the Ensembl genome browser with 121 nucleotide windows. A MATLAB script was used to find CpGs in the Illumina HumanMethylation450 BeadChip (450k) that were located either inside these homolog regions (categorized as “homolog”) or we determined the distance in bp to the center of the closest homolog region (categorized as “distance”). To further investigate age-associated changes at these CpGs the GSE40279 dataset was employed (Hannum et al., 2013). Data was analyzed with the R package GEOquery. Beta-values were used to determine the Pearson correlation with chronological age for all CpGs. The p-value of the correlations (α < 0.01) was estimated with a t-test for linear correlation with Bonferroni-correction for the total amount of CpGs on the 450k array.

### Pyrosequencing

Genomic DNA (about 500ng) was bisulfite converted with the Zymo Research Group EZ DNA Methylation Kit (Zymo Research, Irvine, USA). Primers were designed with PyroMark Assay Design 2.0 software (Metabion, Planegg-Martinsried, Germany; Supplemental Table 1). One of the PCR primers was biotinylated. Age-associated regions were amplified using the PyroMark PCR Kit (Qiagen, Hilden, Germany) with manufactures instructions: 1 μL (approximately 25 ng) of the CT-converted DNA sample was used, with 12.5 μL of the Pyromark MasterMix 2X, 2.5 μL of Coral Load 10X, 0.4 μL of MgCl2, 1.3 μL of forward and reverse primers each and 6 μL of nuclease-free water (25 μL in total per tube). The PCR settings were: initial activation (95°C, 15 min), followed by 50 cycles of denaturation (95°C, 30 sec) + annealing (56°C, 30 sec) + extension (72°C, 30 sec), and one cycle of final extension (72°C, 10 min). Pyrosequencing was then performed on the PyroMark Q48 Autoprep system using the PyroMark Q48 Advanced Reagent Kit according to the manufactures instruction: 10 μL of amplified sample was sequenced with manual primer loading with 2 μL of sequencing primer. The results were analyzed using PyroMark Q48 Advanced software. The pyrosequencing reads covered neighboring CpGs (*Aspa* and region 1: 2 CpGs; *Wnt3a*, *Prima1*, and *Kcns1*; 3 CpGs; *Tbc1d16* and *Hsf4*: 4 CpGs) and their position (pos) was numbered consecutively in the direction of sequencing. For the final selection of a multivariate regression model, the sklearn package in Python was used to identify the combination of 4 CpG sites from different amplicons that minimizes the MAD.

## Results

### Infinium Mouse Methylation analysis

To investigate age-associated DNAm with the Mouse Methylation BeadChip, we used blood samples of 12 B6 mice (11 to 117 weeks old) and 12 DBA mice (6 to 109 weeks old) and we detected that the proportion of methylated CpGs decreases with age (Supplemental Figure 1). Pearson correlation between age and DNAm levels revealed that there was more age-associated hypomethylation than hypermethylation, and this age-associated demethylation was more pronounced in B6 than in DBA mice (Figure 1A). In B6 mice, age-associated hypermethylation was enriched in CpG islands, whereas hypomethylation was rather associated with shore, shelf, and open-sea regions. In contrast, in DBA mice there was rather age-associated hypomethylation at CpG islands, indicating that there are pronounced differences in age-associated DNAm patterns between these mouse strains.

**Figure 1.**
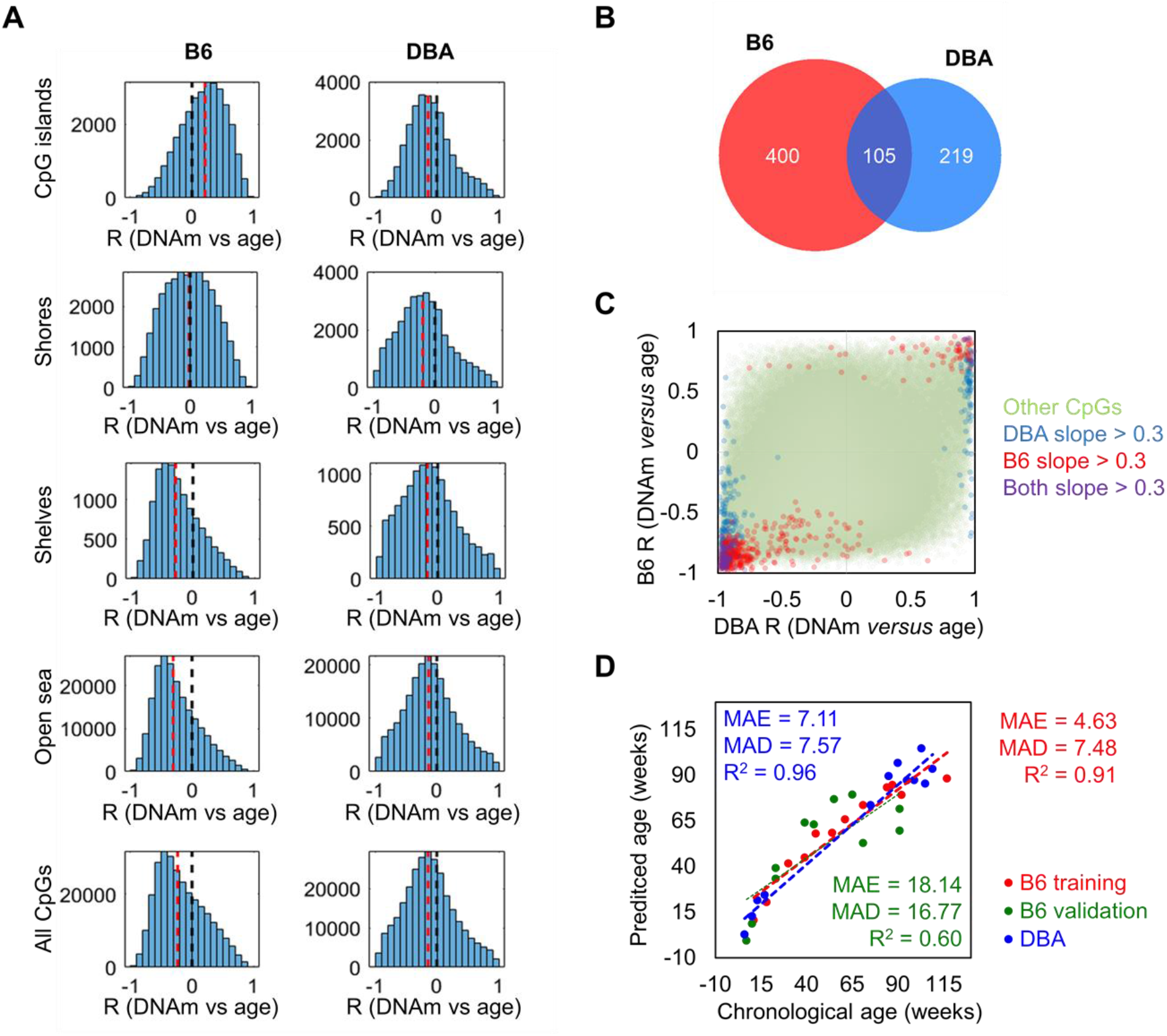
Age-associated DNA methylation differs between mouse strains. **A)** Pearson correlation of DNAm *versus* age (R) for CpGs on the Mouse Methylation BeadChip. Histograms are depicted for different genomic regions (CpG islands, shores, shelves, and open sea) in either B6 or DBA mice. The black dotted line marks no correlation; the red dotted line represents the median of all the CpGs in the respective genomic category. **B)** Venn diagram of the CpGs with slope_CpG_> 0.3 or < −0.3 in linear regressions of age with DNAm. **C)** Comparison of Pearson correlation (R) between age and DNAm in B6 and DBA mice. CpGs that reached the threshold in slope_CpG_ had also high correlations. Notably, several CpGs revealed opposite correlation with age in both mouse strains. **D)** Epigenetic age predictions based on the overlapping 105 age-associated CpGs (MAE = mean average error; MAD = mean absolute deviation; numbers in weeks).

With the aim of selecting candidate CpGs for epigenetic clocks with most prominent changes in DNAm levels in both strains, we filtered on the slope_CpG_ > 0.3 or < −0.3 in linear regression with age: 505 CpGs reached this threshold for B6 mice and 324 CpGs for DBA mice. 105 CpG were in the overlap of both strains (Figure 1B). As anticipated, these CpGs also revealed overall high Pearson correlation with age. When we directly compared the age-associated DNAm between B6 and DBA mice, it became evident that there are marked differences in age-associated DNAm. Several CpGs with positive age-associated DNAm in one strain revealed negative age-associated DNAm in the other strain, and *vice versa* (Figure 1C). Thus, there may even be antagonistic age-associated DNAm changes among mice from distinct strains.

We reasoned that a generally applicable epigenetic clock for mice should focus on the intersection of age-associated DNAm across different mouse strains. Our predictor was therefore trained for B6 mice but based on the 105 overlapping CpGs that also revealed marked age-associated DNAm changes in DBA mice (Figure 1D). The signature based on single-CpG linear predictions was trained on the above mentioned DNAm profiles of the 12 B6 mouse blood samples. The list of 105 CpGs with the slope and intercept for the predictions can be found in Supplemental Table 2. For validation, we analyzed an additional set of 12 B6 mouse blood samples (7 to 91 weeks old). The age predictions correlated with chronological age (R^2^ = 0.60) with a mean average error (MAE) of 18.14 weeks and a mean absolute deviation (MAD) of 16.77 weeks. In analogy, the predictor was applied to the 12 DBA mouse blood samples that were initially considered for the selection of age-associated CpGs but not for training of the model. Despite the different background, the predictor gave relative precise estimates for the DBA cohort (R^2^ = 0.96; MAE = 7.11 weeks; MAD = 7.57 weeks).

### Comparison of DNA methylation profiles in C57BL/6 and DBA/J2 mice

The marked differences in age-related DNAm prompted us to directly compare the methylome of the two mouse strains. Therefore, we first performed a pairwise correlation of all CpGs on the Mouse Methylation BeadChip for each of the individual samples. The DNA methylation profiles of B6 and DBA correlated particularly within the strains in two clear clusters, independent of mouse age, indicating that there are marked differences in their methylome (Figure 2). Furthermore, 17,718 CpGs revealed significant differential methylation (adjusted p-value < 0.01) between B6 and DBA mice. Thus, the methylome of the two inbred mouse strains differs extensively.

**Figure 2.**
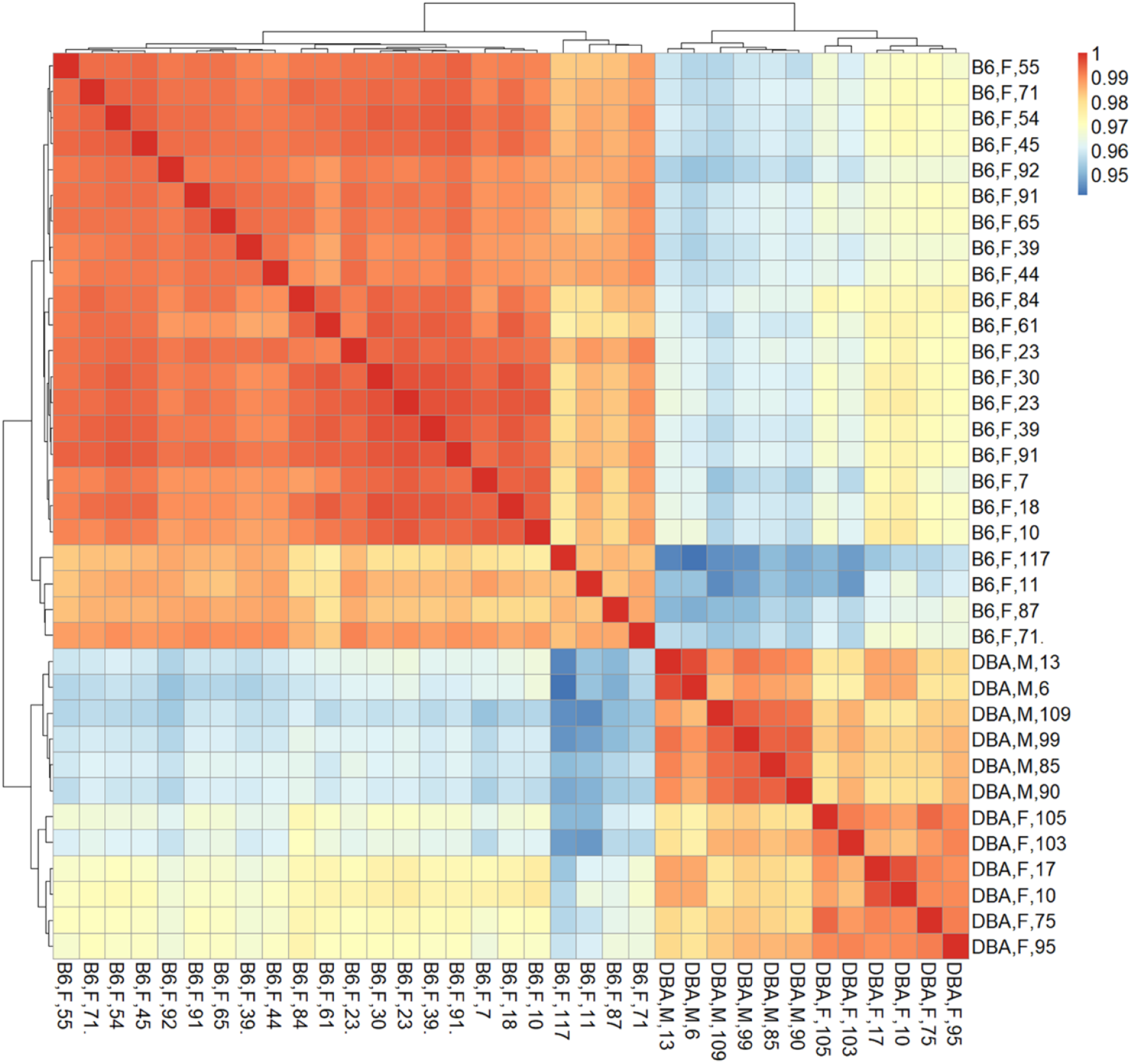
Correlation matrix of DNA methylation profiles. Pearson correlation of DNAm at all CpGs (here also including X and Y chromosome) demonstrated that samples clustered consistently according to the mouse strain, independent of donor age. Sample names indicate mouse strain (B6: C57BL/6; DBA: DBA/2J), sex (M: male, F: female), and age in weeks.

### Tracking of homologous age-related CpG sites between mouse and human

In order to investigate if age-associated DNAm across B6 and DBA mice are also preserved in human, we utilized DNAm profiles of 656 human blood samples for comparison (GSE40279; 19 to 101 years old). When analyzing age-associated CpGs in relation to specific genomic regions we observed enrichment of hypermethylation in CpG islands (Supplemental Figure 2), as described before (Christensen et al., 2009). Homologous regions between human and mice were then identified with *Ensembl*. Out of the 105 overlapping CpGs in B6 and DBA, 83 were located in a region in the mouse genome with a homolog in the human genome. Only 14 CpGs in these homolog regions were also found in the HumanMethylation450 BeadChip, of which seven showed a significant age-methylation correlation in the human homolog region (**Fehler! Verweisquelle konnte nicht gefunden werden.**; Supplemental Table 2). Furthermore, age-associated hyper- and hypomethylation was consistent in both mouse strains and humans for all of the seven selected CpGs. These genomic regions with inter-species conserved age-associated DNA methylation changes were located in the genes aspartoacylase (*Aspa*), WNT family member 3A (*Wnt3a*; cg21934230), heat shock transcription factor 4 (*Hsf4*) and a disintegrin and metalloproteinase with thrombospondin motifs 6 (*Adamts6*).

**Table 1.** Homologous CpGs in mouse and human with high age-methylation correlation.

| CpG ID<br>(Mouse<br>BeadChip) | location in<br>mice (mm10) | Gene | R B6 | R DBA | Homologous location in<br>human (GRCh37) | Homologous<br>CpG Human<br>Methylation<br>450 | R<br>human | p-value<br>human |
| --- | --- | --- | --- | --- | --- | --- | --- | --- |
| cg29748675 | 11:73324349 | *Aspa* | -0.94 | -0.99 | 17:3379534:3379655 | cg02228185 | -0.57 | $4.49 \cdot 10^{-57}$ |
| cg29601161 | 11:59283020 | *Wnt3a* | -0.94 | -0.97 | 1:228202533:228202658 | cg21934230 | -0.35 | $2.83 \cdot 10^{-20}$ |
| cg46095458 | 8:105270995 | *Hsf4* | 0.94 | 0.95 | 16:67199792:67199917 | cg04235075 | 0.25 | $1.43 \cdot 10^{-10}$ |
| cg29601160 | 11:59282958 | *Wnt3a* | -0.91 | -0.98 | 1:228202592:228202736 | cg21934230 | -0.35 | $2.83 \cdot 10^{-20}$ |
| cg29748676 | 11:73324412 | *Aspa* | -0.91 | -0.98 | 17:3379471:3379592 | cg02228185 | -0.57 | $4.49 \cdot 10^{-57}$ |
| cg46095446 | 8:105270822 | *Hsf4* | 0.93 | 0.95 | 16:67199615:67199736 | cg04235075 | 0.25 | $1.43 \cdot 10^{-10}$ |
| cg32016143 | 13:104398027 | *Adamts6* | -0.92 | -0.94 | 5:64558563:64558684 | cg21878650 | -0.36 | $1.21 \cdot 10^{-21}$ |

The homologous genomic regions of *Aspa*, *Wnt3a* and *Hsf4* comprised several neighboring CpGs on the Mouse Methylation BeadChip and the HumanMethylation450 BeadChip. The highest correlated CpGs in these genes were at only few bases apart at the homologous sequences and they revealed similar age-associated hyper- or hypomethylation across species (Figure 3). Notably, the corresponding human region for *ASPA* was already selected as one of three CpGs in our previous aging signature (Weidner et al., 2014). Furthermore, *Hsf4* was one of the three CpGs in our previous aging signature for mice (Han et al., 2018). In addition, two CpGs in the gene proline rich membrane anchor 1 (*Prima1*; cg31044702 and cg29045777) were amongst the top candidates in B6 and DBA (Supplemental Table 2) and this region was also in our previous three CpG signature (Han et al., 2018). Thus, the Mouse Methylation BeadChip identified very similar regions as previous selected from RRBS data (Petkovich et al., 2017), as well as conserved age-associated regions between mice and human.

**Figure 3.**
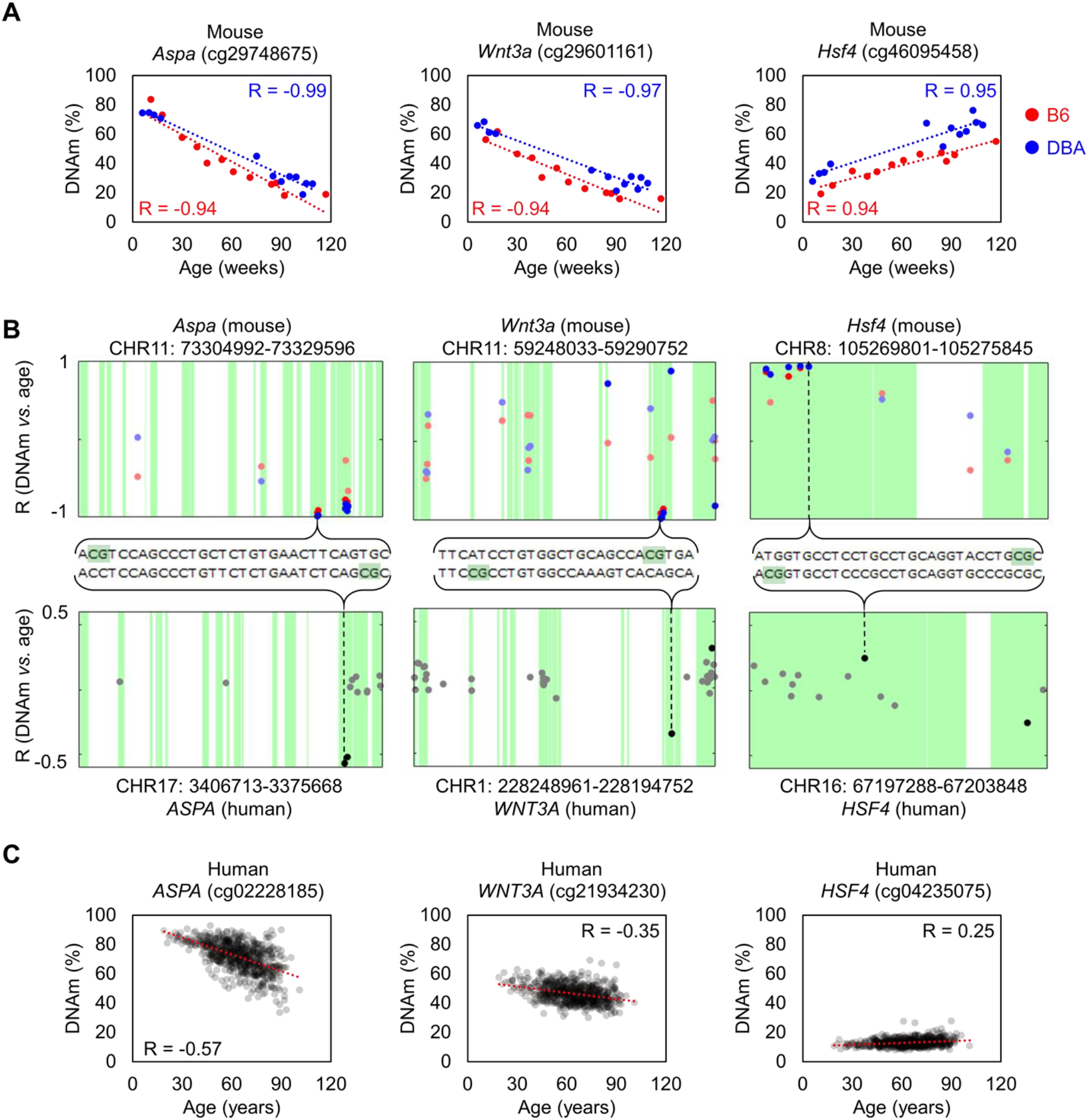
Homologous regions in *Aspa*, *Wnt3a* and *Hsf4* for mice and human. Association of DNAm level (%) and chronological age in the training sets of B6 (n = 12) and DBA (n = 12) mice for the CpG cg29748675 (*Aspa*), cg29748675 (*Wnt3a*), and cg29748675 (*Hsf4*). Genomic location of these CpGs in the Mouse Methylation BeadChip (top) and HumanMethylation450 (bottom). The X-axis indicates the location of each CpG in the sequence of the corresponding gene. For *Aspa* and *Wnt3a*, the horizontal axis for human is inverted (end to start position, from left to right), according to the homology alignment with mouse. Sequences with homology are indicated in green and the Pearson correlation (R) of DNAm with age is presented for each CpG on the BeadChips. CpGs with significant age-associated changes (α = 0.01 after Bonferroni correction) are highlighted with more intense color. **C)** Association of DNAm (%) and age in a dataset of 656 human blood samples (GSE40279; HumanMethylation450 BeadChip).

**Figure 4.**
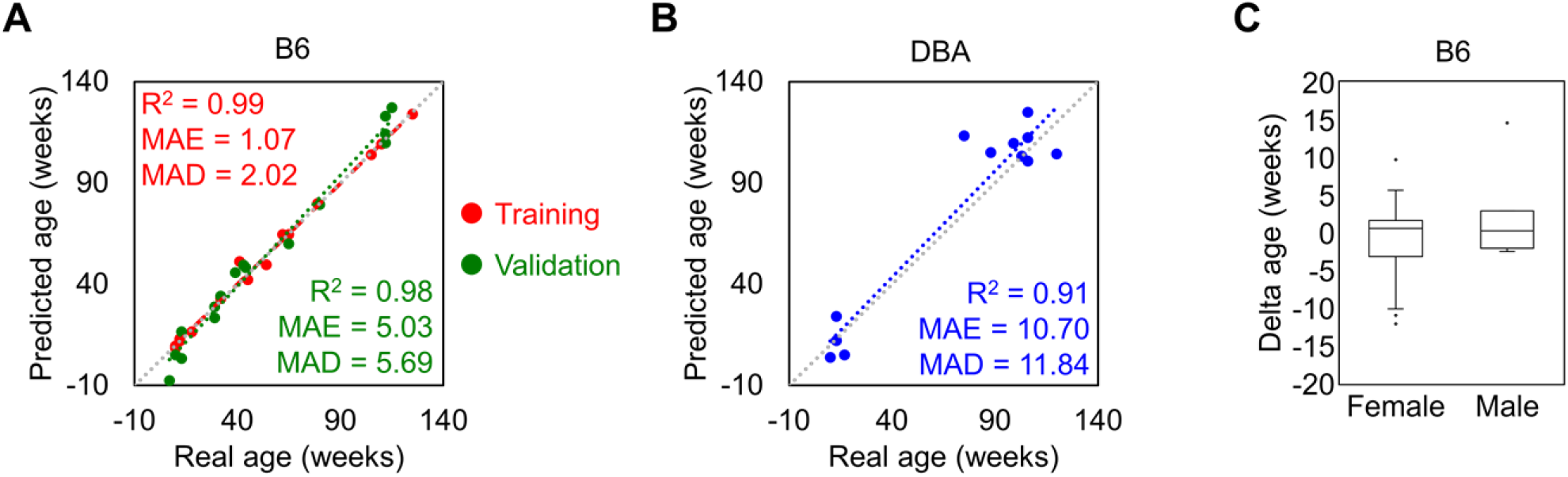
Four CpG epigenetic clock based on pyrosequencing. **A)** Chronological age *versus* predicted age with the 4 CpG model in the B6 training (n = 15) and validation sets (n = 16). **B)** The same 4 CpG predictor was then applied to DBA mice (n = 12). **C)** The deviations of predicted and chronological age (delta age) were plotted for B6 mice (26 female and 5 male). There was no significant difference between male and female samples (p-value = 0.33).

### Pyrosequencing-based age prediction model

Finally, we investigated if the top candidate CpGs that were selected with the Mouse Methylation BeadChip would also provide robust and reliable assays for targeted analysis with pyrosequencing. To this end, the list of 105 CpGs was ranked by mean R^2^ between B6 and DBA mice to then select “Region1” (cg42528232) that is not associated with a specific gene, *Aspa* (cg29748675), and TBC1 Domain Family Member 16 (*Tbc1d16*, cg30271979). Furthermore, the top CpGs with homologies in human were considered (*Aspa*, cg29748675; *Wnt3a*, cg29601161; and *Hsf4,* cg46095458), as well as Potassium Voltage-Gated Channel Subfamily S Member 1 (*Kcns1*), and *Prima1* that were derived from our previous signature (Han et al., 2018). For all these seven regions we established a pyrosequencing assay that comprised a total of 21 CpGs. When we analyzed an independent set of 15 B6 mice (10 to 125 weeks; Supplemental Table 3) all CpGs revealed very high correlation with age (R^2^ >0.8 for each CpG analyzed; Supplemental Figure 3).

To identify the best combination of these age-associated CpGs with regard to MAD of age-predictions, we analyzed every possible multivariate regression model with combinations of 4 CpG sites from different amplicons. The best multivariate model for age-prediction included the variables α (*Aspa* pos1, cg29748675), β (*Hsf4* pos3, 5 bp upstream cg46095458), γ (*Wnt3a* pos2, 9 positions upstream cg29601161) and δ (*Prima1* pos1, cg31044702):

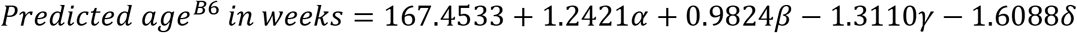

This model was validated with an independent set of B6 mice (age range 7-115 weeks; Supplemental Table 3) and the results revealed very high precision of epigenetic age predictions (R^2^ = 0.98, MAE =5.03 weeks, MAD = 5.69 weeks; **Figure** A). Furthermore, the predictor gave high precision, without clear offset, in DBA mice (n = 12; R^2^ = 0.91, MAE = 10.7 weeks, MAD = 11.84 weeks; **Figure** B). The DNAm levels of the selected CpG sites showed similar association with age in both mouse strains with consistently very high correlations (R^2^ > 0.89) (Supplemental Figure 4). To determine the impact of the sex of mice, we performed a Mann-Whitney U test in the B6 training and validation sets and there was no significant impact of sex on epigenetic age-predictions (p-value = 0.33; **Figure** C).

## Discussion

For development of epigenetic clocks, the Infinium BeadChip technology has proven to be highly efficient for cost-effective integration of multiple datasets. As such platforms were initially not available for non-human species, there have been even attempts to use the Human EPIC BeadChip to investigate potentially conserved CpGs in mice (Gujar et al., 2018). Our exploratory study demonstrates that the Mouse Methylation BeadChip provides reliable insight into age-associated DNAm in mice. It was striking to see that the top candidates derived from RRBS and the microarray analysis (such as *Prima1* and *Hsf4)* did indeed overlap (Han et al., 2018). These novel results thus further substantiate the quality of previous RRBS measurements (Meer et al., 2018; Petkovich et al., 2017).

Our 105 CpG aging signature for the Mouse Methylation BeadChip can be easily adapted by other scientists. It was trained and tested on blood samples and will probably have a big offset in other tissue. These CpGs were selected by the slope of linear regression with age – this criterion favors high absolute DNAm changes with age. We reasoned that this is advantageous for later targeted analysis and to be less susceptible to normalization and inter-study variability. Either way, all selected CpGs had also a high correlation with chronological age, which is commonly applied as filter criterion in other studies (Weidner et al., 2014). In the future, a higher number of measurements will surely enable alternative selection methods, e.g. with elastic net, and multiple variable models. In analogy to human second generation epigenetic clocks, it may even be possible to train murine clocks to capture also other physiological parameters than age alone (Levine et al., 2018). This may provide a better integrative measure of biological age than age-associated DNAm alone.

Larger signatures that comprise hundreds of CpGs may be more robust than targeted assays that only consider one or few CpGs, since they reflect a broader epigenetic pattern (Meer et al., 2018; Thompson et al., 2018; Wagner, 2022). BeadChip technology makes large signatures easily applicable since all relevant CpGs are addressed in each sample. However, adaptation and integration of different microarray datasets remains a major hurdle and age-predictors may become outdated if a BeadChip release is discontinued (Ori et al., 2021). It may therefore be advantageous to rather focus on individual CpGs by targeted methods, such as pyrosequencing, digital droplet PCR or barcoded amplicon sequencing (Wagner, 2022). These methods give very precise and reproducible results on single CpG level and facilitate fast and more cost-effective analysis (Wagner, 2022). Notably, all 21 CpGs covered by our pyrosequencing assay provided very high correlation with age in all training and validation cohorts (R always > 0.8). Our four CpG epigenetic age prediction model (R^2^ = 0.98) thus now outperformes our previously published three CpG signature (R^2^ = 0.95) (Han et al., 2018). Other methods for age prediction in mice have reported lower correlations (R^2^ from 0.64 to 0.91) with a higher number of CpG sites (9 to 582), although their sample size for training was substantially higher (n = 48 to 893) (Simpson and Chandra, 2021). In the future, it will be important to further validate the precision of methylation based age-predictions on larger cohorts.

Our study exemplifies that methylomes of different mouse strains differ extensively. It was striking that in B6 mice age-associated hypermethylation was enriched at CpG islands, whereas in DBA mice CpG islands become rather hypomethylated upon aging. Furthermore, age-associated changes were often divergent in both mouse strains. The DNAm profiles of B6 and DBA clearly clustered together and a substantial fraction of CpGs on the microarray revealed significant differences. This is in line with previous reports on DNAm differences of mouse strains (Gujar et al., 2018; Orozco et al., 2014) and even sub-strains (Sandoval-Sierra et al., 2020). Within the inbred mouse strains there was relatively little epigenetic variation – the inter-individual changes appear to be much less pronounced than in a wild type population. Thus, training of epigenetic clocks within cohorts of one mouse strain, such as B6 and DBA, might correspond to clocks trained on samples that were longitudinally taken from one genetic individual, with much less inter-donor variation. This might also explain, why our signatures provide relevant and highly reproducible results albeit they were only trained on a relatively small set of samples.

It has been suggested that epigenetic clocks are overall accelerated in mice belonging to strains with shorter life spans (Sandoval-Sierra et al., 2020). We have previously demonstrated that our three CpG epigenetic clock was accelerated in DBA as compared to B6, which seemed to reflect the shorter life expectancy (1.87 *versus* 2.42 years) (Han et al., 2018). Moreover, this epigenetic aging clock was accelerated in blood of B6 mice with a congenic DBA version of an age-associated locus on chromosome 11 (Brown et al., 2020). However, the results of our current study indicate that differences in age predictions might also be attributed to the different epigenetic makeups in mouse strains. While some CpGs clearly suggest accelerated epigenetic aging in DBA it may be antagonistic at other sides (e.g. in *Aspa* and *Wnt3a*). For inter-strain comparisons it may therefore be advantageous to consider a wide range of mouse strains to derive an epigenetic clock that is universally applicable across different mouse strains. In addition to various mouse strains also wild type mice should be integrated into such analysis in the future. If such age-associated DNAm changes are even available for multiple different tissues, the signatures might be further trained for an improved multi-tissue mouse aging clock (Thompson et al., 2018). Focusing on conserved age-associated DNAm between murine strains will provide the means to investigate if epigenetic clocks are really accelerated in a specific mouse strain, or if predictions are skewed by the different methylome.

Finally, it is remarkable how some regions exhibit conservation of the age-associated changes in DNAm - not only in different strains, but even across different species. The locus in *Wnt3a* was previously shown to be age-related in human (Brunt et al., 2012; West et al., 2013). Interestingly, the human homolog CpG site *ASPA* (cg02228185) is exactly the same CpG that we selected for our human blood three CpG predictor (Weidner et al., 2014), which was meanwhile used by many other groups (Salameh et al., 2020). The fact that the highest correlated CpGs in the genes *Aspa*, *Wnt3a* and *Hsf4* are within a few bases in mouse-human homology alignment suggests that these specific regions have conserved age-associated methylation patterns across species. In this regard the recently described Mammalian Methylation BeadChip provides a powerful tool (Arneson et al., 2022). It will be interesting to use this platform to better understand how site-specific DNAm changes are conserved across strains and species. What makes these evolutionary conserved regions so special, and if they are mechanistically linked to the aging process needs to further evaluated.

## Author Contributions

J.P.-C. analysed BeadChip data, performed pyrosequencing, and generated the figures. V.T. supported study design and analysis. H.G. provided murine blood samples and supported study design. W.W. designed and supervised the study. J.P.-C. and W.W. wrote the manuscript. All authors have read and approved the final manuscript.

## Acknowledgments

The authors thank Vadim Sakk (Institute of Molecular Medicine, Ulm University) for helping in the collection of mice blood samples.

## References

Arneson, A., Haghani, A., Thompson, M., Pellegrini, M., Kwon, S., Hansen, K., and Hovarth, S. (2022). A mammalian methylation array for profiling methylation levels at conserved sequences. Nature Communications 13, 1–40.

Bell, C.G., Lowe, R., Adams, P.D., Baccarelli, A.A., Beck, S., Bell, J.T., Christensen, B.C., Gladyshev, V.N., Heijmans, B.T., Horvath, S., et al. (2019). DNA methylation aging clocks: challenges and recommendations. Genome Biol 20, 249.

Bibikova, M., Barnes, B., Tsan, C., Ho, V., Klotzle, B., Le, J.M., Delano, D., Zhang, L., Schroth, G.P., Gunderson, K.L., et al. (2011). High density DNA methylation array with single CpG site resolution. Genomics 98, 288–295.

Blueprint-consortium (2016). Quantitative comparison of DNA methylation assays for biomarker development and clinical applications. Nat Biotechnol 34, 726–737.

Bocklandt, S., Lin, W., Sehl, M.E., Sanchez, F.J., Sinsheimer, J.S., Horvath, S., and Vilain, E. (2011). Epigenetic predictor of age. PLoS ONE 6, e14821.

Brown, A., Schuetz, D., Han, Y., Daria, D., Nattamai, K.J., Eiwen, K., Sakk, V., Pospiech, J., Saller, T., van Zant, G., et al. (2020). The lifespan quantitative trait locus gene Securin controls hematopoietic progenitor cell function. Haematologica 105, 317–324.

Brunt, K.R., Zhang, Y., Mihic, A., Li, M., Li, S.H., Xue, P., Zhang, W., Basmaji, S., Tsang, K., Weisel, R.D., et al. (2012). Role of WNT/beta-catenin signaling in rejuvenating myogenic differentiation of aged mesenchymal stem cells from cardiac patients. Am J Pathol 181, 2067–2078.

Christensen, B.C., Houseman, E.A., Marsit, C.J., Zheng, S., Wrensch, M.R., Wiemels, J.L., Nelson, H.H., Karagas, M.R., Padbury, J.F., Bueno, R., et al. (2009). Aging and environmental exposures alter tissue-specific DNA methylation dependent upon CpG island context. PLoS Genet 5, e1000602.

Fraga, M., Ballestar, E., Paz, M., Ropero, S., Serien, F., Ballestar, M., Cigudosa, J., Benitez, J., and Esteller, M. (2005). Epigenetic differences arise during the lifetime of monozygotic twins. PNAS 102, 10604–10609.

Gujar, H., Liang, J.W., Wong, N.C., and Mozhui, K. (2018). Profiling DNA methylation differences between inbred mouse strains on the Illumina Human Infinium MethylationEPIC microarray. PLoS One 13, e0193496.

Han, Y., Eipel, M., Franzen, J., Sakk, V., Dethmers-Ausema, B., Yndriago, L., Izeta, A., de Haan, G., Geiger, H., and Wagner, W. (2018). Epigenetic age-predictor for mice based on three CpG sites. Elife 7.

Han, Y., Nikolic, M., Gobs, M., Franzen, J., de Haan, G., Geiger, H., and Wagner, W. (2020). Targeted methods for epigenetic age predictions in mice. Sci Rep 10, 22439.

Hannum, G., Guinney, J., Zhao, L., Zhang, L., Hughes, G., Sadda, S., Klotzle, B., Bibikova, M., Fan, J.B., Gao, Y., et al. (2013). Genome-wide Methylation Profiles Reveal Quantitative Views of Human Aging Rates. Mol Cell 49, 459–367.

Horvath, S. (2013). DNA methylation age of human tissues and cell types. Genome Biology 14, 1–19.

Koch, C.M., and Wagner, W. (2011). Epigenetic-aging-signature to determine age in different tissues. Aging (Albany NY) 3, 1018–1027.

Levine, M.E., Lu, A.T., Quach, A., Chen, B.H., Assimes, T.L., Bandinelli, S., Hou, L., Baccarelli, A.A., Stewart, J.D., Li, Y., et al. (2018). An epigenetic biomarker of aging for lifespan and healthspan. Aging (Albany NY) 10, 573–591.

Marioni, R.E., Shah, S., McRae, A.F., Chen, B.H., Colicino, E., Harris, S.E., Gibson, J., Henders, A.K., Redmond, P., Cox, S.R., et al. (2015). DNA methylation age of blood predicts all-cause mortality in later life. Genome Biol 16, 25.

Meer, M.V., Podolskiy, D.I., Tyshkovskiy, A., and Gladyshev, V.N. (2018). A whole lifespan mouse multi-tissue DNA methylation clock. Elife 7.

Niu, L., Xu, Z., and Taylor, J.A. (2016). RCP: a novel probe design bias correction method for Illumina Methylation BeadChip. Bioinformatics (Oxford, England) 32, 2659–2663.

Ori, A.P.S., Horvath, S., and Ophoff, R.A. (2021). A systematic evaluation of 41 DNA methylation predictors across 101 data preprocessing and normalization strategies highlights considerable variation in algorithm performance. BioRxiv Preprint.

Orozco, L.D., Rubbi, L., Martin, L.J., Fang, F., Hormozdiari, F., Che, N., Smith, A.D., Lusis, A.J., and Pellegrini, M. (2014). Intergenerational genomic DNA methylation patterns in mouse hybrid strains. Genome Biol 15, R68.

Petkovich, D., Podolskiy, D., Lobanov, A., Lee, S., Miller, R., and Gladyshev, V. (2017). Using DNA Methylation Profiling to Evaluate Biological Age and Longevity Interventions. Cell Metabolism 25, 954–960.

Salameh, Y., Bejaoui, Y., and El Hajj, N. (2020). DNA Methylation Biomarkers in Aging and Age-Related Diseases. Front Genet 11, 171.

Sandoval-Sierra, J.V., Helbing, A.H.B., Williams, E.G., Ashbrook, D.G., Roy, S., Williams, R.W., and Mozhui, K. (2020). Body weight and high-fat diet are associated with epigenetic aging in female members of the BXD murine family. Aging Cell 19, e13207.

Simpson, D.J., and Chandra, T. (2021). Epigenetic age prediction. Aging Cell 20, e13452.

Stubbs, T., Bonder, M., Stark, A., Krueger, F., von Meyenn, F., Stegle, O., and Reik, W. (2017). Multi-tissue DNA methylation age predictor in mouse. Genome Biology 18, 1–14.

Thompson, M.J., Chwialkowska, K., Rubbi, L., Lusis, A.J., Davis, R.C., Srivastava, A., Korstanje, R., Churchill, G.A., Horvath, S., and Pellegrini, M. (2018). A multi-tissue full lifespan epigenetic clock for mice. Aging (Albany NY) 10, 2832–2854.

Wagner, W. (2017). Epigenetic aging clocks in mice and men. Genome Biol 18, 107.

Wagner, W. (2022). How to Translate DNA Methylation Biomarkers Into Clinical Practice. Front Cell Dev Biol 10, 854797.

Wang, T., Tsui, B., Kreisberg, J.F., Robertson, N.A., Gross, A.M., Yu, M.K., Carter, H., Brown-Borg, H.M., Adams, P.D., and Ideker, T. (2017). Epigenetic aging signatures in mice livers are slowed by dwarfism, calorie restriction and rapamycin treatment. Genome Biol 18, 57.

Weidner, C.I., Lin, Q., Koch, C.M., Eisele, L., Beier, F., Ziegler, P., Bauerschlag, D.O., Jockel, K.H., Erbel, R., Muhleisen, T.W., et al. (2014). Aging of blood can be tracked by DNA methylation changes at just three CpG sites. Genome Biol 15, R24.

West, J., Beck, S., Wang, X., and Teschendorff, A.E. (2013). An integrative network algorithm identifies age-associated differential methylation interactome hotspots targeting stem-cell differentiation pathways. Scientific Reports 3, 1630.

Xu, Z., Niu, L., Li, L., and Taylor, J. (2016). ENmix: a novel background correction method for Illumina HumanMethylation450 BeadChip. Nucleic Acids Research 44, 1–6.

Yuan, R., Tsaih, S.W., Petkova, S.B., Marin de Evsikova, C., Xing, S., Marion, M.A., Bogue, M.A., Mills, K.D., Peters, L.L., Bult, C.J., et al. (2009). Aging in inbred strains of mice: study design and interim report on median lifespans and circulating IGF1 levels. Aging Cell 8, 277–287.

Zhou, L., Ng, H.K., Drautz-Moses, D.I., Schuster, S.C., Beck, S., Kim, C., Chambers, J.C., and Loh, M. (2019). Systematic evaluation of library preparation methods and sequencing platforms for high-throughput whole genome bisulfite sequencing. Sci Rep 9, 10383.

