## Supplemental Figures and Table S1 for "Epigenetic clocks for mice based on age-associated regions that are conserved between mouse strains and human"

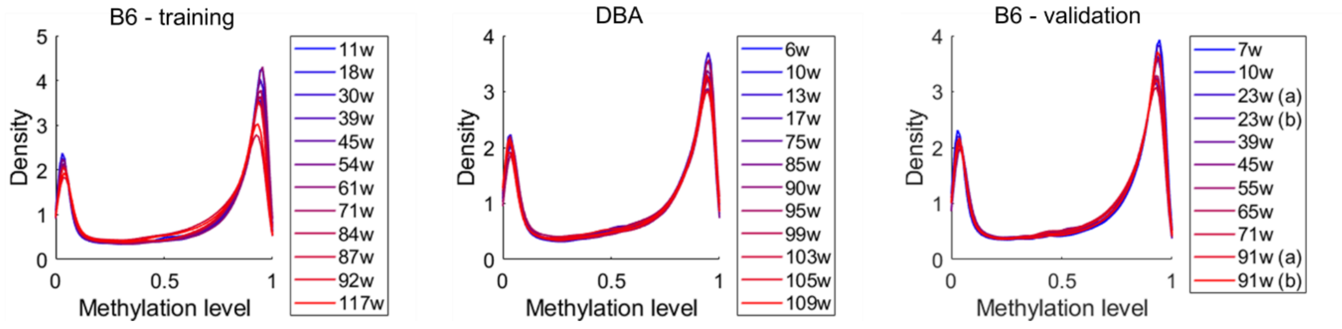


**Supplemental Figure 1. Methylation density plots.**

Histograms demonstrate the DNA methylation distribution of CpGs in the B6 training, DBA, and B6 validation cohort. Most CpGs reveal either low or high DNAm levels (close to 0% or 100%). The color code reflects age in weeks (blue = young; red = old) and demonstrates hypomethylation with age.

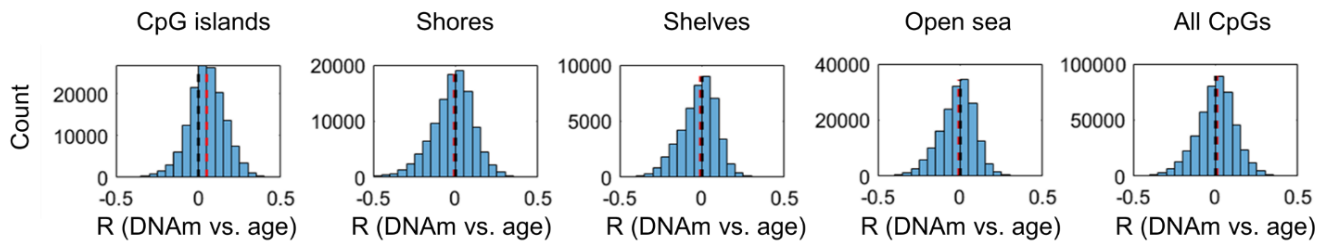


**Supplemental Figure 2. Pearson correlation of DNAm *versus* chronological age in human.**

Histograms show the distribution of Pearson correlation (R) in the Infinium HumanMethylation450 BeadChip datasets (Hannum et al., 2013) (GSE40279) in analogy to Figure 1A. The black dotted lines represent no correlation; the red dotted lines represents the median of the correlation of all CpGs in the respective genomic category. In humans, there is a moderate tendency that age-associated gains in methylation are enriched at CpG islands.

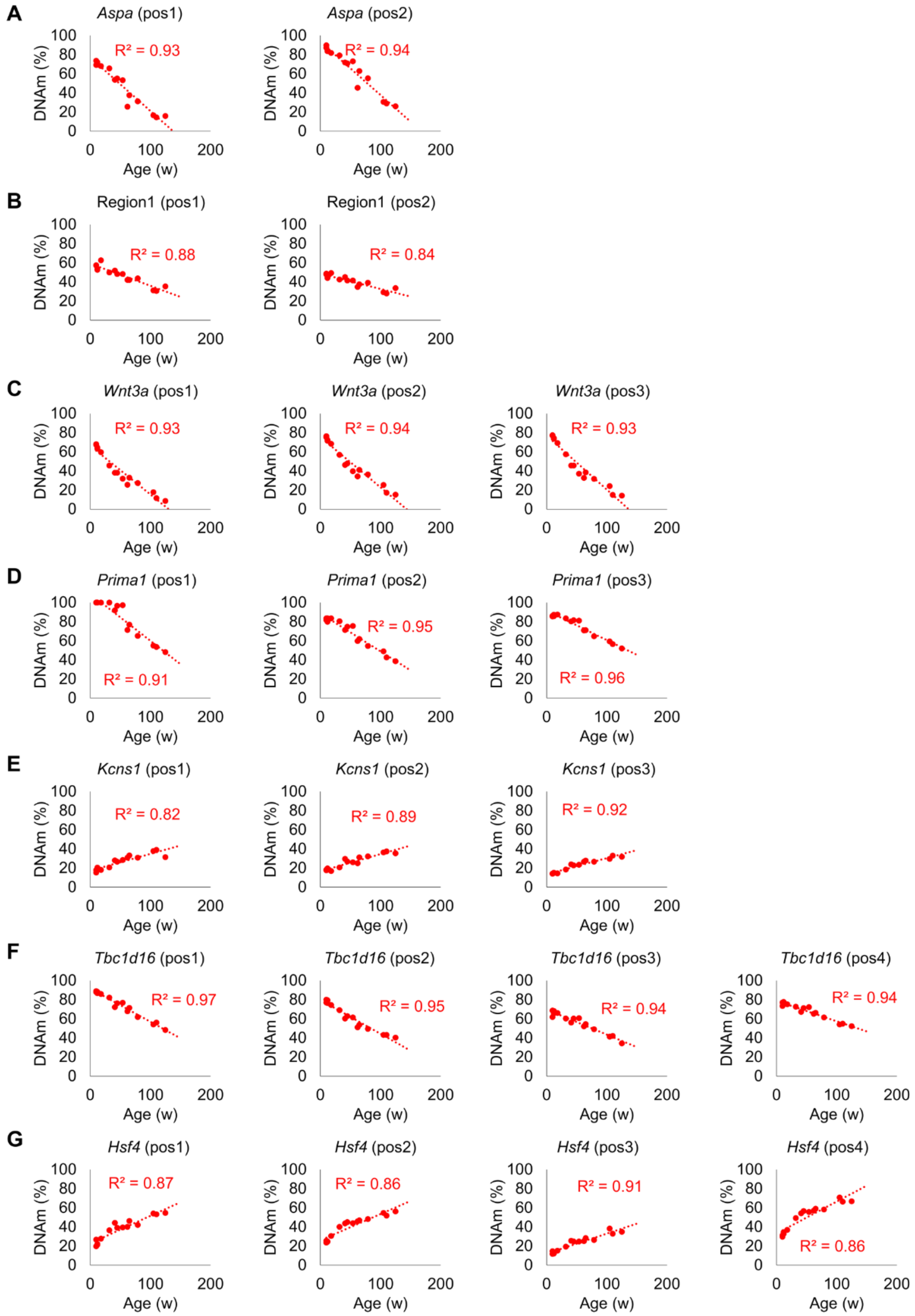


**Supplemental Figure 3. Pyrosequencing results for the B6 training set.**

Pyrosequencing was performed for seven different amplicons covering a total of 21 different CpGs. The graphs present the age in weeks *versus* the methylation percentage of the CpG sites in **A)** *Aspa* **B)** Region1 **C)** *Wnt3a* **D)** *Prima1* **E)** *Kcns1* **F)** *Tbc1d16*, and **G)** *Hsf4*. The CpGs of the selected amplicons revealed overall very high correlations with 14 out of 21 CpGs showing $R^2 > 0.9$, and all of them showing $R^2 > 0.8$.

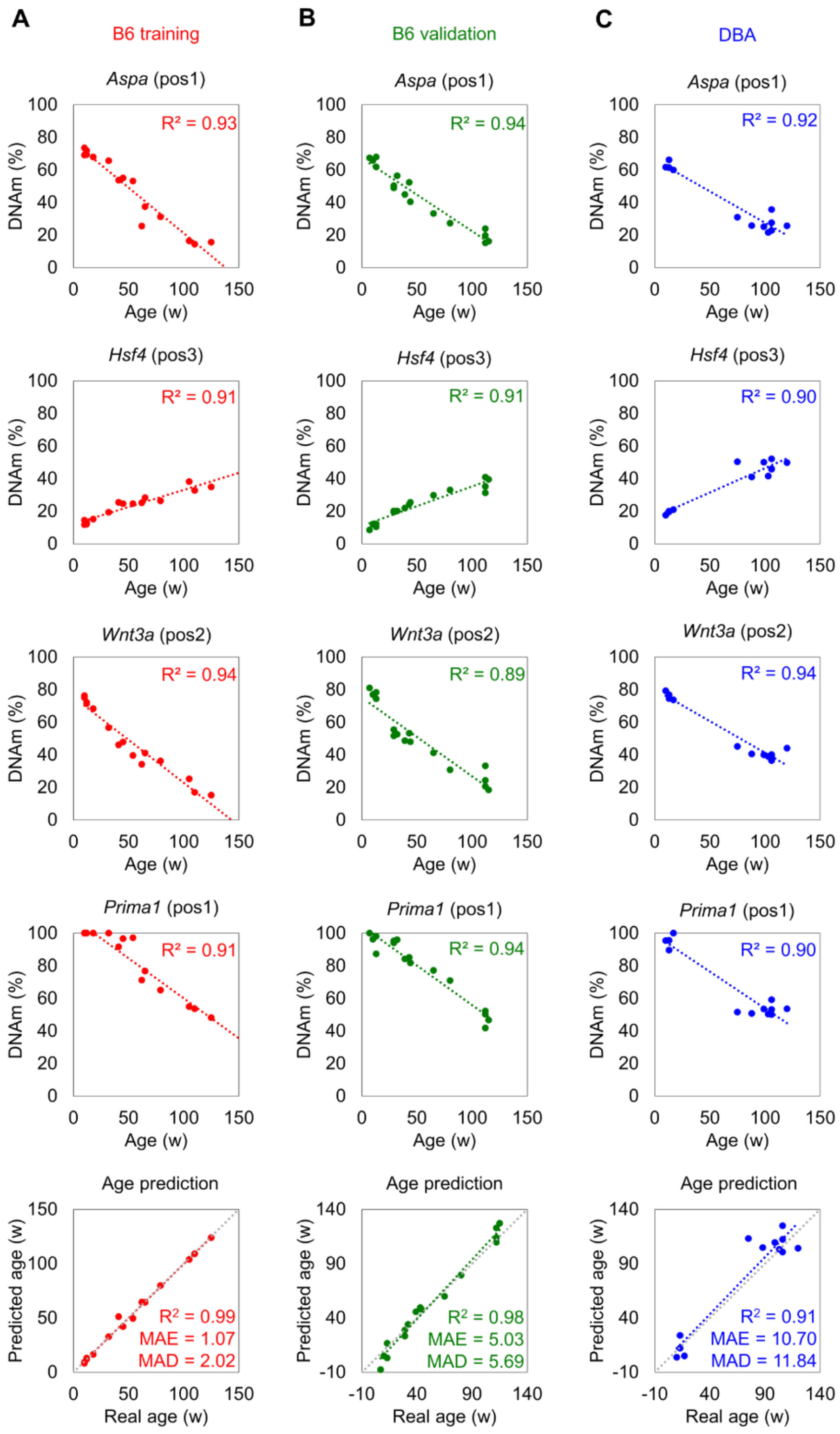


**Supplemental Figure 4. Pyrosequencing results for the 4 CpGs in training and validation sets.**

The graphs present the age in weeks *versus* the methylation percentage of the CpG sites in **A)** B6 training set, **B)** B6 validation set, and **C)** DBA validation set. All the selected CpG sites showed a very high correlation with age ($R^2 > 0.89$) in all the different cohorts.

**Supplemental table 1. List of primers for pyrosequencing.**

| Primer | Sequence |
|---|---|
| ***Prima1:*** | |
| Forward | 5'-TTGTGTTTAATTAGGAGAGGTAAATTATGAATTAGGTTTATA-3' |
| Reverse | 5'-Biotin-CAAAATTAATTACACCAACTTATAACCTACTATTC-3' |
| Sequencing | 5'-AATTATGAATTAGGTTTATATTT-3' |
| ***Hsf4:*** | |
| Forward | 5'-GTGAGTAGTAAGGTGGGATAAATTGTAGAAAAAATG-3' |
| Reverse | 5'-Biotin-TCCCTACTCTCCTACACTCCTCTCAAAACTTA-3' |
| Sequencing | 5'-ATTGTAGAAAAAATGGGAA-3' |
| ***Kcns1:*** | |
| Forward | 5'-GGTTGAGAGGGTGGTAGAAGAAGTTG-3' |
| Reverse | 5'-Biotin-ACTCCCCTCCATCCCTACCATATACATCCA-3' |
| Sequencing | 5'-GAAGATATTTAGAAGTTGAATT-3' |
| ***Aspa:*** | |
| Forward | 5‘-TTTTTGGATTGGTAATGAATGG-3‘ |
| Reverse | 5‘-Biotin-AAAAAACCTATTAAAAAAATTACCATCTT-3‘ |
| Sequencing | 5‘-GGATTGGTAATGAATGGTT-3‘ |
| ***Tbc1d16:*** | |
| Forward | 5‘-AATTGTGGAGTAATGTTAGTGG-3‘ |
| Reverse | 5‘-Biotin-ATAACTTTTAAATCCTCCTATCACA-3‘ |
| Sequencing | 5‘-GTGGTAGGTTGGGTT-3‘ |
| ***Wnt3a:*** | |
| Forward | 5‘-Biotin-GAAGGAGGAGGGAGAAAAATTATT-3‘ |
| Reverse | 5‘-CCCCATCCTTAAACTAAAAATTTCATC-3‘ |
| Sequencing | 5‘-ACTAAAAATTTCATCCTATAACTAC-3‘ |
| ***CHR5:105409216*** | **(Region1):** |
| Forward | 5‘-TAGAAGTTAAGGTTTAAGAAAATAGTGT-3‘ |
| Reverse | 5‘-Biotin-AAACATATTCAAACCACCAAAACTA-3‘ |
| Sequencing | 5‘-AGTGTTTAGAAGTTTAGTTAG-3‘ |

**Supplemental table 2. List of 105 age-associated CpGs in B6 and DBA.**

The table provides CpG IDs on the Mouse Methylation BeadChip, chromosome location, information of associated genes and correlation (R and $R^2$) in B6 and DBA, human homolog regions, correlation of human CpGs if available (R and $R^2$), categorization if the relevant CpG is directly within the homologous sequence (HOM.) or distant (DIST.). For each individual CpG the intersection and slope is predicted for a single CpG predictor based on the blood samples of the B6 training set. This table is provided as separated EXCEL file.

**Supplemental table 3. Pyrosequencing results.**

The table provides all pyrosequencing results for the training and validation sets, for all seven amplicons and each CpG within these amplicons. This table is provided as separated EXCEL file.
